# art_modern: An Accelerated ART Simulator of Diverse Next-Generation Sequencing Reads

**DOI:** 10.64898/2026.02.20.707060

**Authors:** Zhejian Yu

**Affiliations:** Centre of Biomedical Systems and Informatics of Zhejiang University-University of Edinburgh Institute, Zhejiang University School of Medicine, Hangzhou 310003, China; Department of Bioengineering, University of Pennsylvania, Philadelphia, PA 19104, USA; Center for Computational and Genomic Medicine, Children’s Hospital of Philadelphia, Philadelphia, PA 19104, USA

## Abstract

**Summary:** Fast simulation of next-generation sequencing (NGS) data is vital for software development and benchmarking. Here we describe art_modern, an accelerated ART simulator that can simulate various NGS data. We accelerated ART using updated sampling algorithms, single-instruction multiple-data (SIMD) instruction-set extensions (ISEs), thread- and node-level parallelism, and an asynchronous output writer, while enabling simulation of transcriptome profiling data by supporting contig-specific coverage with strand information. The new implementation was benchmarked against popular performance-oriented NGS simulators, revealing a 75–77% reduction in CPU time and a 15–24 times acceleration in wall-clock time on a multi-core machine compared to the original implementation. With this simulator, the process of developing and benchmarking NGS sequence analysis algorithms can be largely accelerated.

**Availability and Implementation:** The software is implemented in C++17 with CMake as the building system. It can be built and executed on a modern GNU/Linux operating system with Boost, Zlib, and a C++17 compiler, with further acceleration available using Intel OneAPI C++/DPC++ compilers and Intel oneAPI MKL random generators. The software is available at https://github.com/YU-Zhejian/art_modern under the GNU General Public License v3.

**Contact:** Zhejian Yu

## Introduction

High-performance simulation of realistic next-generation sequencing (NGS) reads could facilitate algorithm development and benchmarking. Primarily developed for the 1000 Genome Project, Artificial Read Transcription (ART) is a popular NGS simulator that has been cited more than 1,000 times in Scopus (2026/1/13) [1]. In a recent benchmark, it outperformed most of its competitors in aspects of speed and the realism of quality scores [2]. Here we introduce art_modern, a modern re-implementation of the ART simulator with enhanced performance and additional functionality. In addition to all the features of the original ART, our software supports thread-based parallelism, accelerated random number generators, synthesis of BAM files, and different sequencing depths for individual contigs. Additionally, it includes a faster profiler generator that generates ART-compatible error profiles from FASTQ/SAM/BAM/NCBI SRA input. We believe that this simulator can largely accelerate the testing and benchmarking of NGS-related bioinformatics algorithms.

## Methods

### Algorithm Overview

The simulation algorithm of the art_modern simulator is an accelerated re-implementation of the classic ART algorithm, which mainly includes 4 steps: (1) Parsing arguments and submit parallelized simulation executors to the job pool; (2) In the executors, a DNA insert will be extracted and normalized from the provided input, with indels, qualities, and single-nucleotide substitutions added based on the given ART error profile and controlling parameters; (3) Generating pairwise alignments (PWAs) and append them to thread-level dispatchers; (4) Dispatchers convert the PWAs to final on-disk data structure and submits them to low-level output queues, which writes them to disk using various supported output formats (Figure 1).

**Figure 1.**
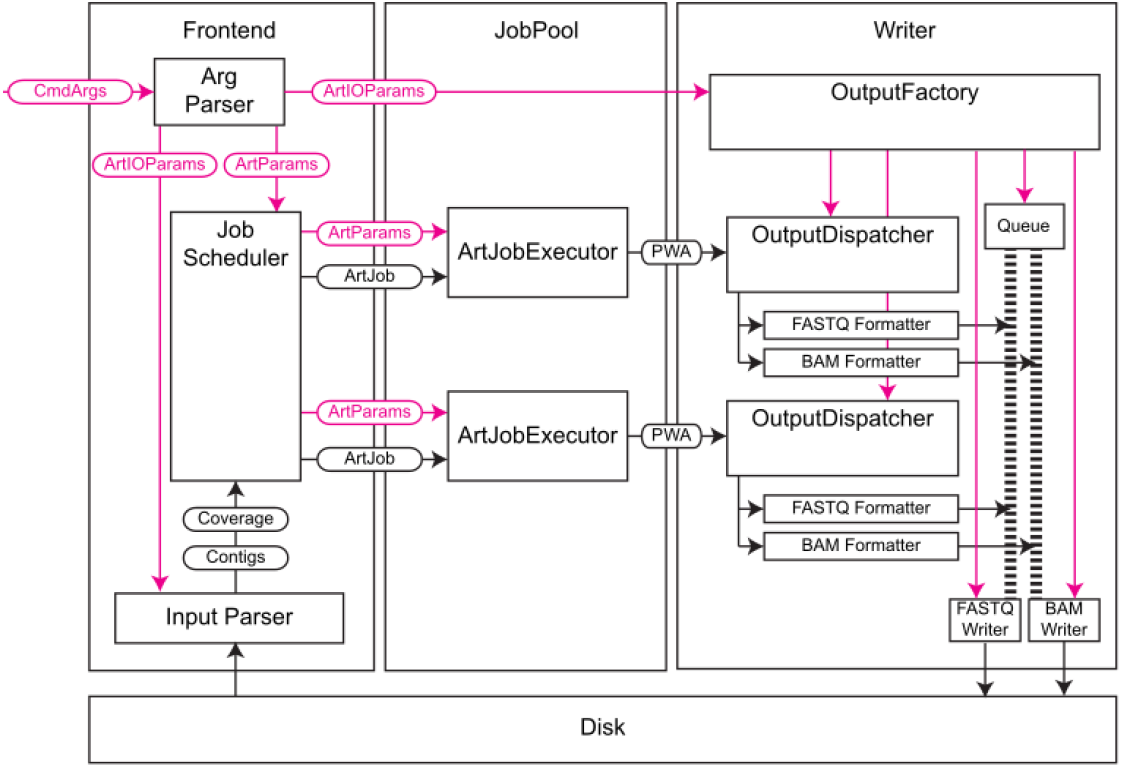
Architecture overview of art_modern.

In addition to whole-genome sequencing (WGS) supported by the original ART, our simulator also supports transcriptome and template (amplicon) sequencing. While the WGS mode supports only unified coverage (same coverage across all contigs), transcriptomics and template mode support different coverage information for each contig and strand. This allows simulating differentially expressed genes or isoforms from gene expression profiles generated by an upstream simulator. All 3 modes support single-end, pair-end (where read 1 and read 2 are facing the center of the insert), and mate-pair (where read 1 and read 2 are facing the terminals of the insert) simulation. For the WGS and transcriptome modes, the simulator will try to mimic ultrasound fragmentation of the input library. Specifically, in single-end mode, the simulated reads will start at random positions along the contig, ensuring a uniform distribution. In pair-end and mate-pair simulations, the insert length is first generated from a Gaussian (Normal) distribution using user-provided parameters. The insert position will then be drawn from a uniform distribution with reads generated from both sides of the insert. For the template mode, the simulator will only generate sequencing reads from the first base of the input template (c)DNAs under single-end and take the provided template as the entire insert under pair-end or mate-pair mode. The generated insert sequence will then be normalized by transforming lower-case nucleotides to uppercase and assigning nucleotides other than A, T, C, and G to N.

In the second step, random insertions and deletions will be introduced at random positions of the read using a fixed insertion and deletion rate provided by the user. The quality value for each position will then be generated from a known error profile that records quality values and their frequencies for each position. The original ART used a binary search over accumulated probabilities with complexity as high as O(log(n)). In this work, we improved the efficiency of discrete distribution sampling using Walker’s algorithm [3], which reduces this complexity to O(1) with negligible overhead in memory. With this approach, the overall simulation speed is largely accelerated. The substitutions will ultimately be generated based on the simulated quality scores.

The generated reads will then be transformed into PWA, a data structure that contains read name, course contig and position, reference sequence, generated read, generated quality, and base-to-base alignment information. The generated PWAs will then be passed to the writers of dedicated formats, as detailed in the next section.

Accelerations of multiple utility functions, like sequence reversals, are done through single-instruction multiple-data (SIMD) instruction set extensions (ISEs) that are present in modern x86_64 CPUs [4]. With such technologies, we can achieve significant performance improvements by grouping similar operations into SIMD-accelerated function calls. The program also supports acceleration using alternative malloc/free implementations like jemalloc [5], mi-malloc [6], or tcmalloc [7].

### Details on Input Routines

Multiple input parsers were implemented to accommodate the wide range of formats and simulation strategies acceptable by the simulator. Like other simulators, the widely used FASTA format is supported by all simulation modes, with additional support over the PBSIM3 Transcripts format for its transcriptome or template mode [8].

Recent advances in long-read sequencing and assembly algorithms enable the assembly of enormous genomes and the annotation of complex transcriptomes. The simulation on those genomes with a relatively small footprint is addressed in 2 ways in this simulator: For the WGS simulation mode, we introduce HTSLib for parsing large genomes in FASTA format with exceptionally large contigs [9]. For transcriptome and template simulation mode, the simulator supports progressive reading of a theoretically infinite stream of short (c)DNA molecules in small batches, and generates the simulated data as a continuous stream. This makes the simulator able to accept (c)DNA fragments from an upstream simulator that mimics the library construction processes to introduce Illumina sequencing noise.

### Details on Random Number Generation and Parallelism

Most of the acceleration is done using faster randomization libraries and algorithms. In ART, the rand function family from the C standard libraries and routines from the GNU Science Library (GSL) were interchangeably used, which largely complicates the software engineering tasks, generates inconsistencies between different platforms, while adding GSL as a required dependency [10]. In this work, we unified the choice of random number generators using the Mersenne Twister pseudo-random number generator (mt19937) as it is well-defined, widely adopted, and has a long period, enough for simulation purposes [11]. The simulator supports the mt19937 algorithm implemented in STL, Intel oneAPI Math Kernel Library (oneMKL)^1^, and Boost::Random. Under Intel platforms, the oneMKL implementation is considerably faster due to its SIMD-accelerated bulk generation of random numbers. However, this may not hold across other platforms, and benchmarks are required to determine the most suitable algorithm for each platform. Another pseudo-random number generator, the Permuted Congruential Generators (PCG) family of random generator, were also supported as a high-performance alternative for the STL mt19937 random generator if other acceleration libraries are not available [12]. The mt19937 implementation in GSL is not used due to performance issues.

A thread-based parallelism is implemented to support multi-core CPUs. Jobs containing a job ID, a reference sequence fetcher, and coverage information will be generated in the main thread and passed to the corresponding job executors, which will generate all reads with the additional ART- and NGS-specific parameters it holds. Each thread will hold one executor instance with parallelism realized using the thread pool implemented using either Boost::ASIO or BS::thread_pool [13]. A fail-safe mode without any parallelism was also implemented.

Different simulation modes and input parsers yield distinct parallelism strategies. For WGS simulation, each thread gets either a pointer to a shared in-memory reference sequence list (if the reference is read into the memory) or owns a wrapper of faidx_t* (if the reference is parsed using HTSLib), with coverage information being divided and assigned to each thread. For transcriptome and template simulation, however, each thread receives only a portion of the reference sequences with full coverage information, which is small and easier to duplicate. It is worth noting that if the reference sequence had already been read into memory, it would need to be copied for each thread, resulting in a memory footprint 2 times the size of the reference genome.

For output, all generated PWAs will be passed to an output dispatcher, which triggers registered writers that convert them to the native in-memory datagram of the corresponding output format. The converted instances would finally be added in parallel to fixed-sized multi-producer single-consumer Moody Camel concurrent queues of unique pointers of their native formats held by each writer [14], [15]. With such, the writers can accept alignments in parallel without the cost of mutexes. Writers of the Sequence Alignment/Map (SAM) and Binary Alignment/Map (BAM) formats are supported through HTSLib, which adds another check for possibly malformed alignments, SAM tags, and parallel compression [16]. A special headless SAM/BAM format is designed to accommodate the “stream” input parser. Compared to the common SAM/BAM format, this format does not include contig information in its header, as alignment information is stored in the OA tag of each alignment record. With these improvements, the simulator only needs to traverse the input reference stream once, generating continuous outputs that facilitate streamlining processing with UNIX pipelines. Other output formats, such as FASTA and FASTQ, are also supported. It is worth noting that for pair-end and mate-pair simulations, FASTAs and FASTQs are generated in an unordered, interleaved format, with manual separation and sorting required before feeding them to most aligners.

The potential of multi-node high-performance clusters (HPCs) can also be utilized using the Message Passing Interface (MPI)-enabled version of the simulator [17]. With MPI, the simulator can be launched simultaneously across multiple computing nodes, fully exhausting the cluster’s computational power. After being launched, individual simulation processes will parse arguments on their own, processing the input file with total depth divided by number of MPI processes (under WGS mode) or by skipping contigs based on rank and number of MPI processes (under other modes), launching multiple threads as indicated by parallelism parameters, while generating output files that are suffixed with the MPI rank it is assigned to. The logging module is also engineered so that only the MPI process with rank 0 reports to standard error, with logs from other MPI processes written to files, preventing interference with the HPC log systems. A side effect of this approach is that the generated output files should be manually merged after the simulator finishes. Support for special files such as UNIX pipelines, I/O redirections, and FIFOs is also dropped in MPI-enabled art_modern.

Finally, this work enables parallelized, reproducible read simulation. While other simulators, such as pIRS, only allow the seed to be set when the simulator is running with 1 process [18], this work used a master-seed strategy to achieve this. After the simulator starts, a master seed will be generated and distributed to all MPI processes if MPI is enabled. Each process will then offset the seed by MPI rank, and use it to seed a PCG32 random generator. The seeds generated by the PCG random generator will then be used to seed all jobs, thus ensuring reproducibility. The current limitation is that reproducibility can only be achieved when the seed, the reference, and the parallelization strategy (i.e., number of MPI processes and threads) are defined.

### Utilities

The original ART is shipped with a Perl-implemented profile generator, and this study implemented an accelerated one in C++ with parallelization and the ability to parse SAM/BAM/CRAM and NCBI Short Read Archive (SRA) files using HTSLib and NCBI C++ Toolkit. Despite its speed, the new profile generator also fixed a bug in the original one that would offset the quality values by 1. This tool is released with art_modern. We also implemented a Python tool for synthesizing a FASTQC-like report from an ART profile and a SAM/BAM validator, available at https://github.com/YU-Zhejian/art_modern_utils.

### Benchmarks

art_modern 1.3.3, compiled using GCC and Intel compiler, is benchmarked against the original ART 2016.06.05, DWGSIM 0.1.15 [19], pIRS 2.0.0, and the wgsim simulator bundled with SAMtools 1.21 [16], [20]. Although wgsim and DWGSIM generate only dumb quality scores, we still include them because they are the fastest NGS simulators in benchmarks. All tools, except art_modern (GCC), were compiled with the Intel OneAPI DPC++/C++ Compiler 2025.3.2.20260112 using the -march=native, -mtune=native, and -O3 compiler options. wgsim and art_modern were linked to HTSLib version 1.23.0, compiled with the same compiler and flags, and the original ART was linked to GSL 2.8, compiled with the same compiler and flags. art_modern (GCC) is compiled with the GNU Compiler Collection (GCC) 13.3.0-6ubuntu2∼24.04, using PCG as the random number generator. On the other hand, art_modern (Intel) is compiled with the Intel compiler and Intel oneMKL 2025.0.3.0 for the random number generator. Both art_modern were linked to {fmt} 12.1.0, Boost 1.83.0, and HTSLib 1.23. Other tools involve GNU Time 1.9, seqkit 2.9.0 [21], seqtk 1.2 [22], R 4.5.2 [23], dplyr 1.1.4 [24], and ggplot2 3.4.4 [25]. The benchmarks were executed on an HP ZBook Power 15.6-inch G10 Mobile Workstation PC with an Intel i7-13700H CPU, 32 GiB of DDR4 memory, the Linux Mint 22.2 (based on Ubuntu 24.04 LTS) operating system, and the Linux 6.8.0-100-generic kernel.

To ensure fair measurement, PE100 and PE300 error profiles are trained for ART, art_modern, and pIRS using datasets from CNR0028307 (Illumina HiSeq2000) [26] and SRR16074289 (Illumina HiSeq2500) [27]. The FASTQs were used to train ART and art_modern quality profiles using the art_modern profile builder mentioned above. They’re then respectively aligned to the reference genome of *E. coli* (NCBI RefSeq GCF_000005845.2) and *G. max* (NCBI RefSeq GCF_000004515.6) using Burrows-Wheeler aligner (BWA) 0.7.18-r1243 [28], [29] to train the pIRS Base-Calling profile, and only primary alignments were used to train the pIRS indel profiles. Since ART does not support GC bias or profile-based Indel simulation, those features were disabled during pIRS execution. The wgsim/DWGSIM simulation of variants was also disabled, and DWGSIM was modified to suppress output compression. The benchmark code is provided in https://github.com/YU-Zhejian/art_modern_benchmark_other_simulators.

For memory and temporal complexity under different sequencing depths, we generated PE100 and PE300 data using *C. elegans* reference genome (ce11) from UCSC (hereby referred to as the “genome” data) and the first 2,500 mRNA sequences of *H. Sapiens* reference genome (hg38) that contain no ambiguous bases with its length exceeding 1000 bp from UCSC (hereby referred to as the “transcriptome” data) with a gradient sequencing depth of 1×, 2×, 4×, 8×, and 16×, with art_modern and pIRS running using 6 threads. The CPU time of art_modern in both datasets is significantly lower than that of the original ART and lies between wgsim and DEGSIM, with its wall-clock time lower than all other simulators, thanks to parallelism (Figure 2A). However, art_modern consumes more memory than other simulators. A possible explanation is that other simulators run simulations on a single contig at a time, whereas art_modern reads a batch of contigs into memory before simulating to support parallelization.

**Figure 2.**
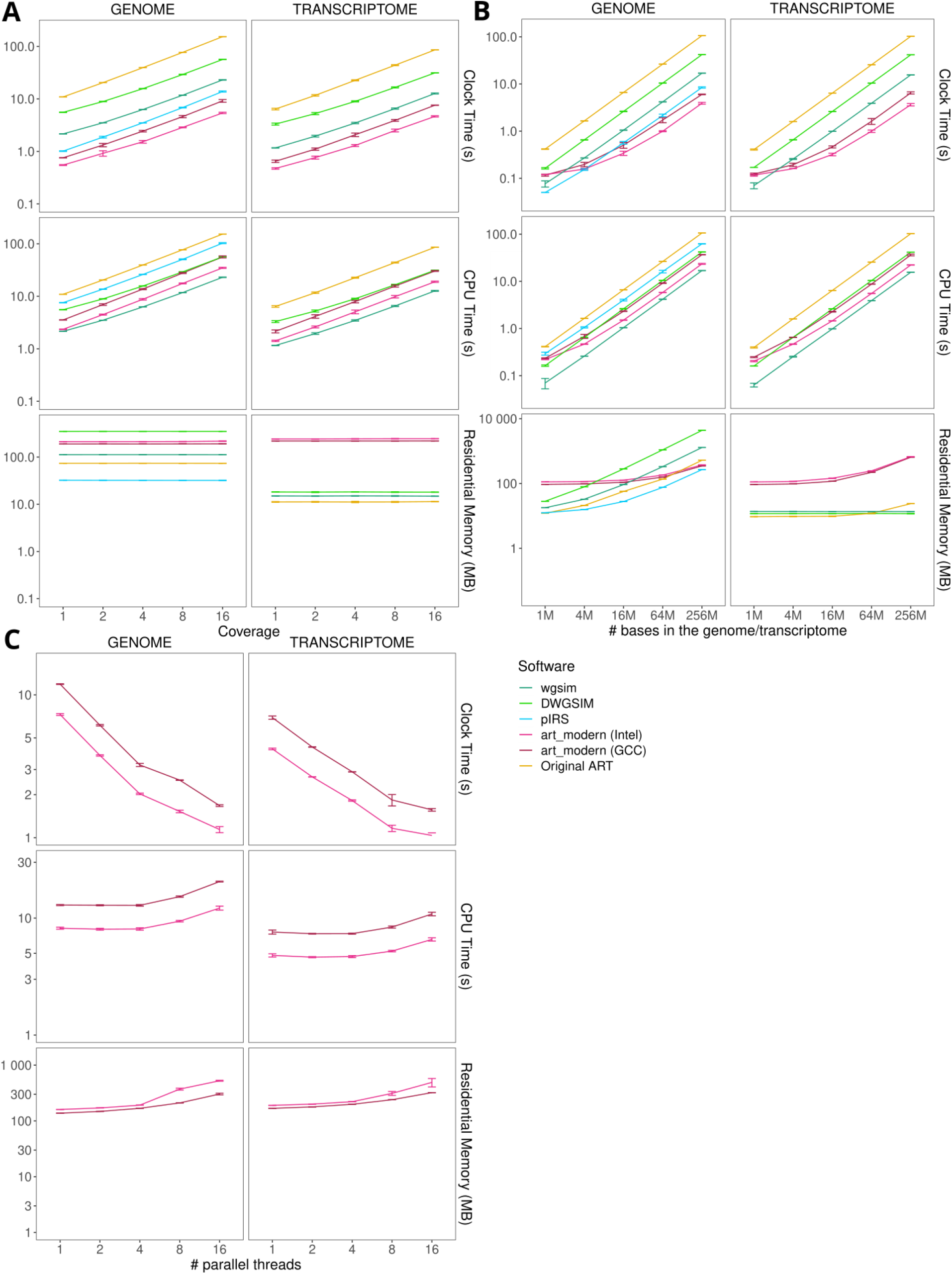
Benchmark Results under PE100 Condition. **A:** Wall-clock time, CPU time, and residential memory size of each program on the genome and transcriptome dataset of different sequencing depths. The CPU time of pIRS in the transcriptome dataset is exceptionally high (1200–1500 seconds), so excluded. **B:** Wall-clock time, CPU time, and resident memory size of each program on the genome and transcriptome of different sizes. C: Wall-clock time, CPU time, and residential memory size of art_modern (GCC) and art_modern (Intel) under different numbers of threads.

To evaluate performance across varying reference genomes and transcriptome sizes, we generated several single-contig simulated genomes ranging from 1 Mbp to 256 Mbp, along with multiple simulated transcriptomes containing 1k to 256k transcripts, each approximately 1 kbp in length. Each simulator simulates raw reads under a depth of 4×. From the results, we observe that art_modern is slower than wgsim/DWGSIM on small reference genomes/transcriptomes but faster on larger inputs (Figure 2B). This may be due to the overhead of preparing quality score generators, as art_modern outperforms pIRS and the original ART across all genome sizes.

The parallelization strategy of art_modern is also tested in the benchmark, where art_modern was built using GCC and Intel compilers, generating data at 4× sequencing depth for the *C. Elegans* genome and the *H. Sapiens* long transcriptome. From the results, we observe that clock time decays with increasing thread count, while CPU time and memory consumption steadily increase due to parallelization overhead (Figure 2C). It can also be observed that the Intel build of art_modern generally consumes more memory than the GCC build, which may be due to the internal random number cache used by the MKL random generator, which effectively speeds up random number generation by producing them in bulk.

## Discussion and Conclusion

To conclude, in this work, we enhanced the original ART with bug fixes, significant performance improvements, greater scalability, and richer functionality, at the cost of increased memory consumption. The resulting software can be a drop-in replacement for the original ART and applied to scenarios such as benchmarking DNA- or RNA-Seq alignment algorithms, testing the performance of self-built (sc)RNA-Seq pipelines, or pressure-testing omics analysis pipelines on a cluster. This simulator is best suited for GNU/Linux-based high-end desktops (HEDTs) with multiple cores and a fast solid-state drive (SSD). It may also work on GNU/Linux laptops, HPCs, or Apple macOS/FreeBSD workstations. The acceleration strategy and data structures involved can be applied to a wider range of simulators, including those targeting long-read sequencing. Further acceleration using GPU-based random number generators is possible, but is not implemented in this version due to the additional complexities in the installation and maintenance processes.

The software is freely available on GitHub (https://github.com/YU-Zhejian/art_modern) with documentation (https://yu-zhejian.github.io/art_modern_docs/index.html), benchmarks (https://github.com/YU-Zhejian/art_modern_benchmark_other_simulators), and utilities (https://github.com/YU-Zhejian/art_modern_utils) under the GNU General Public License (GPL) v3, the same as the original ART. It is also published on BioConda (https://bioconda.github.io/recipes/art_modern/README.html) with an MPI-enabled variant (https://bioconda.github.io/recipes/art_modern-openmpi/README.html), with utilities published in PyPI (https://pypi.org/project/art-modern-utils/).

## Supporting information

Raw Data for Figures and R-Produced Figures

## Acknowledgments

The author would like to express their sincere gratitude to the original authors of ART and HTSLib for their wonderful software.

## Supplementary Files

The raw figures and data files for the benchmark are uploaded as a ZIP file.

## Footnotes

1 Note that on Intel oneMKL, we actually use SFMT19937, which is an MT19937 variant that allows further acceleration using SIMD technologies.

## References

[1] W. Huang, L. Li, J. R. Myers, and G. T. Marth, “ART: a next-generation sequencing read simulator.,” Bioinformatics (Oxford, England), vol. 28, no. 4, pp. 593–594, Feb. 2012, doi: 10.1093/bioinformatics/btr708.

[2] M. Milhaven and S. P. Pfeifer, “Performance evaluation of six popular short-read simulators,” Heredity, vol. 130, no. 2, pp. 55–63, Dec. 2022, doi: 10.1038/s41437-022-00577-3.

[3] A. J. Walker, “An Efficient Method for Generating Discrete Random Variables with General Distributions,” ACM Trans. Math. Softw., vol. 3, no. 3, pp. 253–256, Sep. 1977, doi: 10.1145/355744.355749.

[4] Intel Corporation, “Intel® 64 and IA-32 Architectures Software Developer’s Manual,” vol. 1, Sep. 2016, [Online]. Available: https://software.intel.com/content/dam/www/public/us/en/documents/manuals/64-ia-32-architectures-software-developer-vol-1-manual.pdf

[5] J. Evans, “A Scalable Concurrent malloc(3) Implementation for FreeBSD,” Apr. 2006, [Online]. Available: https://people.freebsd.org/~jasone/jemalloc/bsdcan2006/jemalloc.pdf

[6] D. Leijen, B. Zorn, and L. De Moura, “Mimalloc: Free List Sharding in Action,” in Programming Languages and Systems, vol. 11893, A. W. Lin, Ed., in Lecture Notes in Computer Science, vol. 11893., Cham: Springer International Publishing, 2019, pp. 244–265. doi: 10.1007/978-3-030-34175-6_13.

[7] Z. Zhou et al., “Characterizing a Memory Allocator at Warehouse Scale,” in Proceedings of the 29th ACM International Conference on Architectural Support for Programming Languages and Operating Systems, Volume 3, La Jolla CA USA: ACM, Apr. 2024, pp. 192–206. doi: 10.1145/3620666.3651350.

[8] Y. Ono, M. Hamada, and K. Asai, “PBSIM3: a simulator for all types of PacBio and ONT long reads,” NAR Genomics and Bioinformatics, vol. 4, no. 4, p. lqac092, Dec. 2022, doi: 10.1093/nargab/lqac092.

[9] J. K. Bonfield et al., “HTSlib: C library for reading/writing high-throughput sequencing data,” GigaScience, vol. 10, no. 2, p. giab007, Jan. 2021, doi: 10.1093/gigascience/giab007.

[10] M. Galassi et al., GNU scientific library reference manual: For GSL version 1.12, 3. ed. Bristol: Network Theory, 2009.

[11] M. Matsumoto and T. Nishimura, “Mersenne twister: a 623-dimensionally equidistributed uniform pseudo-random number generator,” ACM Transactions on Modeling and Computer Simulation, vol. 8, no. 1, pp. 3–30, Jan. 1998, doi: 10.1145/272991.272995.

[12] M. E. O’Neill, “PCG: A Family of Simple Fast Space-Efficient Statistically Good Algorithms for Random Number Generation,” Harvey Mudd College, Claremont, CA, HMC-CS-2014-0905, Sep. 2014. [Online]. Available: https://www.cs.hmc.edu/tr/hmc-cs-2014-0905.pdf

[13] B. Shoshany, “A C++17 Thread Pool for High-Performance Scientific Computing,” SoftwareX, vol. 26, p. 101687, 2024, doi: 10.1016/j.softx.2024.101687.

[14] C. Desrochers, “A Fast General Purpose Lock-Free Queue for C++.” Nov. 2014. [Online]. Available: https://moodycamel.com/blog/2014/a-fast-general-purpose-lock-free-queue-for-c++

[15] C. Desrochers, “Detailed Design of a Lock-Free Queue.” Nov. 2014. [Online]. Available: https://moodycamel.com/blog/2014/detailed-design-of-a-lock-free-queue

[16] H. Li et al., “The Sequence Alignment/Map format and SAMtools,” Bioinformatics, vol. 25, no. 16, Art. no. 16, Aug. 2009, doi: 10.1093/bioinformatics/btp352.

[17] The MPI Forum, “MPI: A message passing interface,” in Supercomputing ‘93:Proceedings of the 1993 ACM/IEEE Conference on Supercomputing, Nov. 1993, pp. 878–883. doi: 10.1145/169627.169855.

[18] X. Hu et al., “pIRS: Profile-based Illumina pair-end reads simulator.,” Bioinformatics (Oxford, England), vol. 28, no. 11, pp. 1533–1535, Jun. 2012, doi: 10.1093/bioinformatics/bts187.

[19] N. Homer, DWGSIM. (2011). [Online]. Available: https://github.com/nh13/DWGSIM

[20] P. Danecek et al., “Twelve years of SAMtools and BCFtools,” GigaScience, vol. 10, no. 2, p. giab008, Jan. 2021, doi: 10.1093/gigascience/giab008.

[21] W. Shen, S. Le, Y. Li, and F. Hu, “SeqKit: A Cross-Platform and Ultrafast Toolkit for FASTA/Q File Manipulation,” PLOS ONE, vol. 11, no. 10, p. e0163962, Oct. 2016, doi: 10.1371/journal.pone.0163962.

[22] H. Li, seqtk. (2014). [Online]. Available: https://github.com/lh3/seqtk

[23] R Core Team, R: A Language and Environment for Statistical Computing. Vienna, Austria: R Foundation for Statistical Computing, 2024. [Online]. Available: https://www.R-project.org/

[24] H. Wickham, R. François, L. Henry, K. Müller, and D. Vaughan, dplyr: A Grammar of Data Manipulation. 2023. [Online]. Available: https://dplyr.tidyverse.org

[25] H. Wickham, ggplot2: Elegant Graphics for Data Analysis. Springer International Publishing, 2016. doi: 10.1007/978-3-319-24277-4.

[26] T. Hu et al., “Comparison of the DNBSEQ platform and Illumina HiSeq 2000 for bacterial genome assembly,” Sci Rep, vol. 14, no. 1, p. 1292, Jan. 2024, doi: 10.1038/s41598-024-51725-0.

[27] H. Liu et al., “Structural variation of mitochondrial genomes sheds light on evolutionary history of soybeans,” The Plant Journal, vol. 108, no. 5, pp. 1456–1472, 2021, doi: 10.1111/tpj.15522.

[28] H. Li and R. Durbin, “Fast and accurate short read alignment with Burrows-Wheeler transform,” Bioinformatics, vol. 25, no. 14, Art. no. 14, Jul. 2009, doi: 10.1093/bioinformatics/btp324.

[29] H. Li and R. Durbin, “Fast and accurate long-read alignment with Burrows-Wheeler transform,” Bioinformatics, vol. 26, no. 5, Art. no. 5, Mar. 2010, doi: 10.1093/bioinformatics/btp698.

