## Supplementary figures and images for "art_modern: An Accelerated ART Simulator of Diverse Next-Generation Sequencing Reads"

### time_memory-size_100.pdf

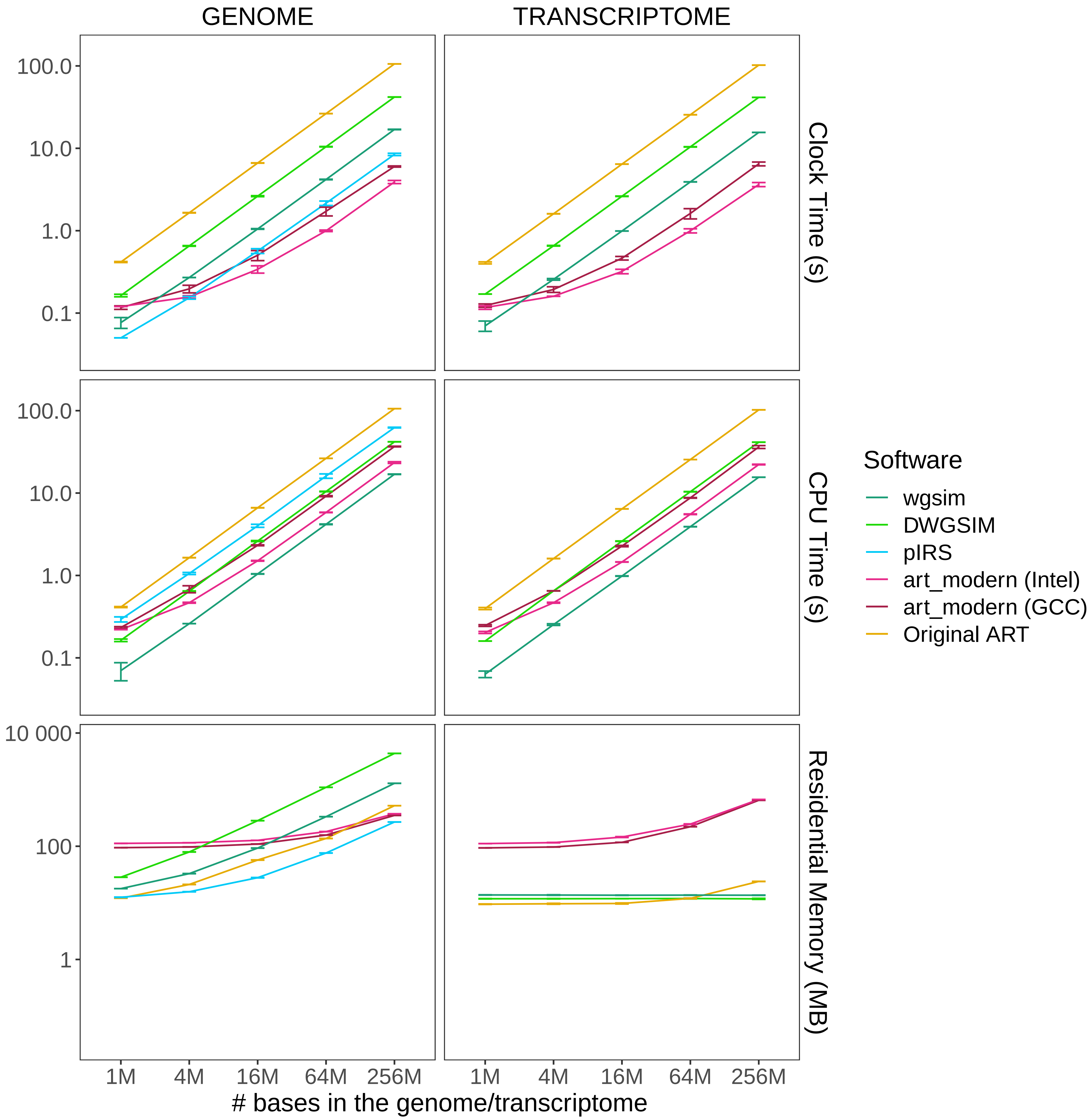

### time_memory-size_300.pdf

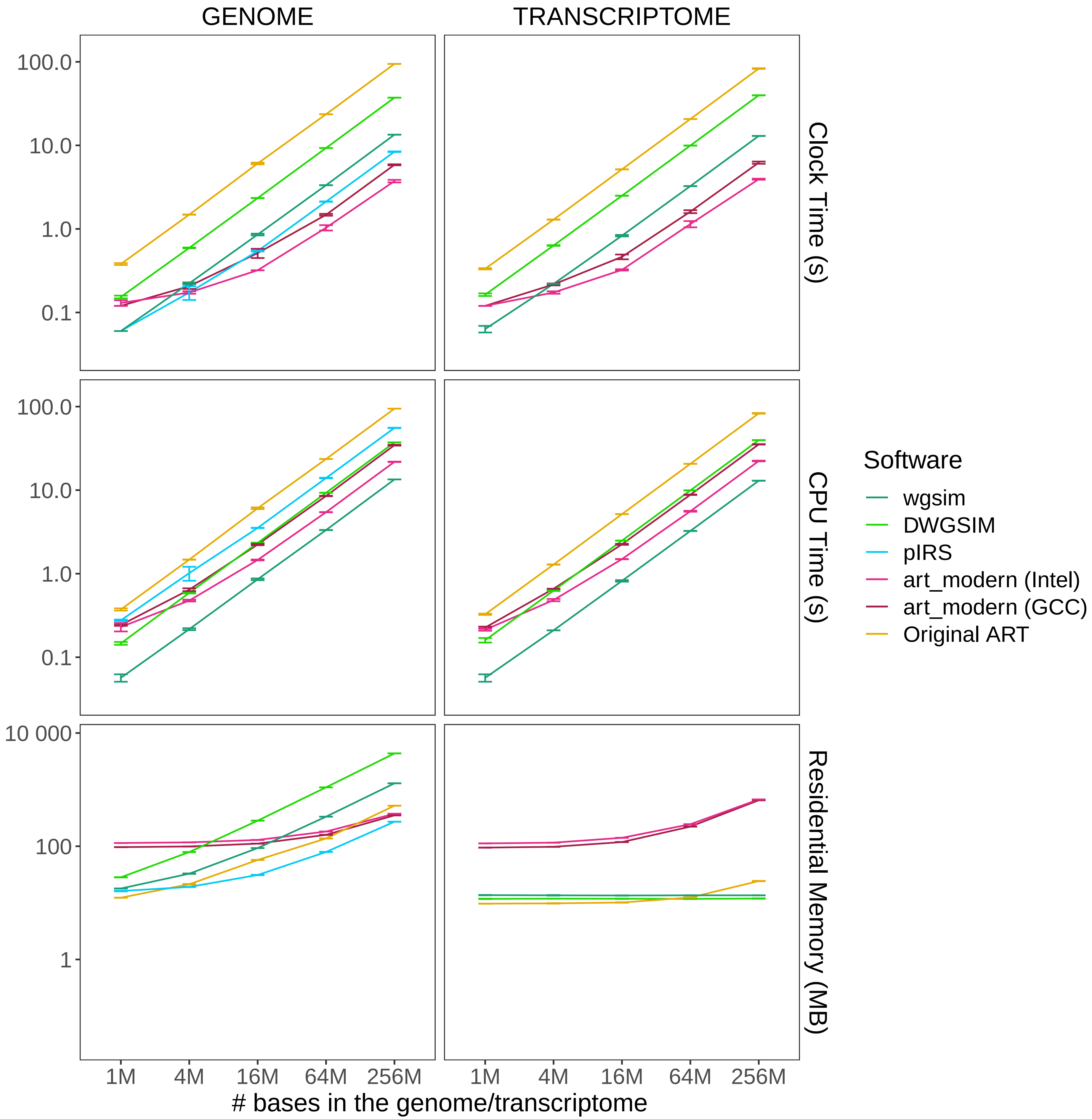

### time_memory-speedup_100.pdf

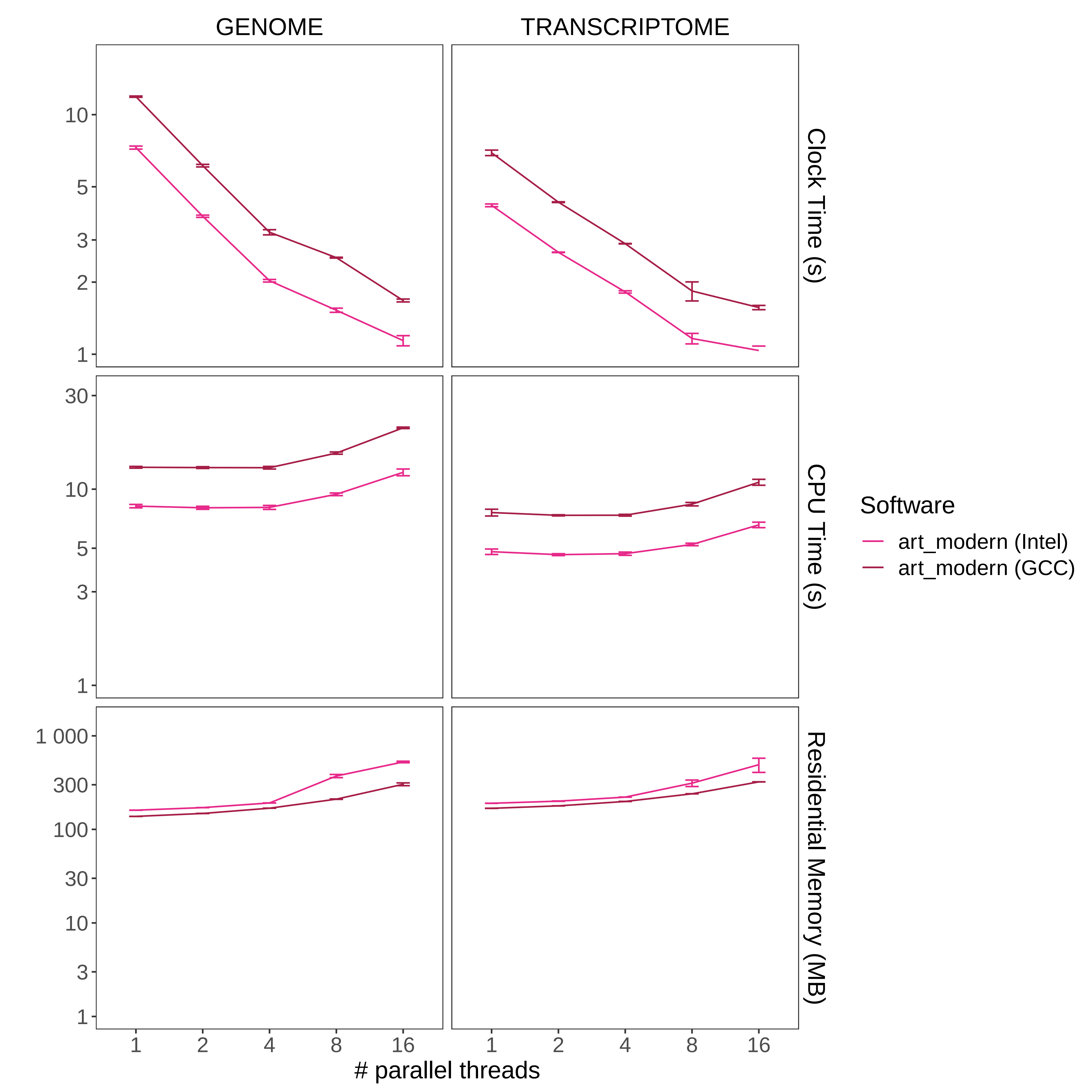

### time_memory-speedup_300.pdf

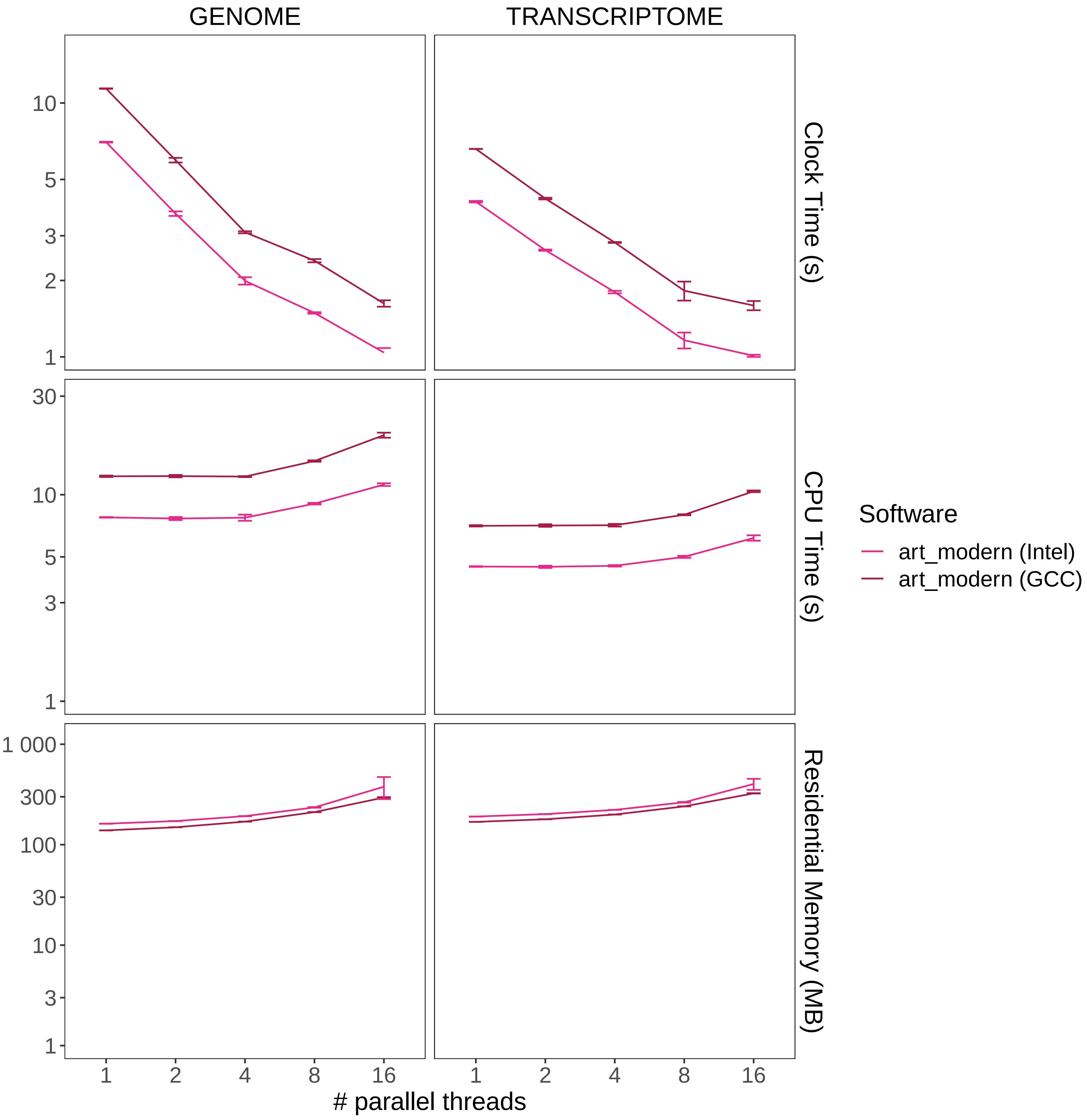

### time_memory_100.pdf

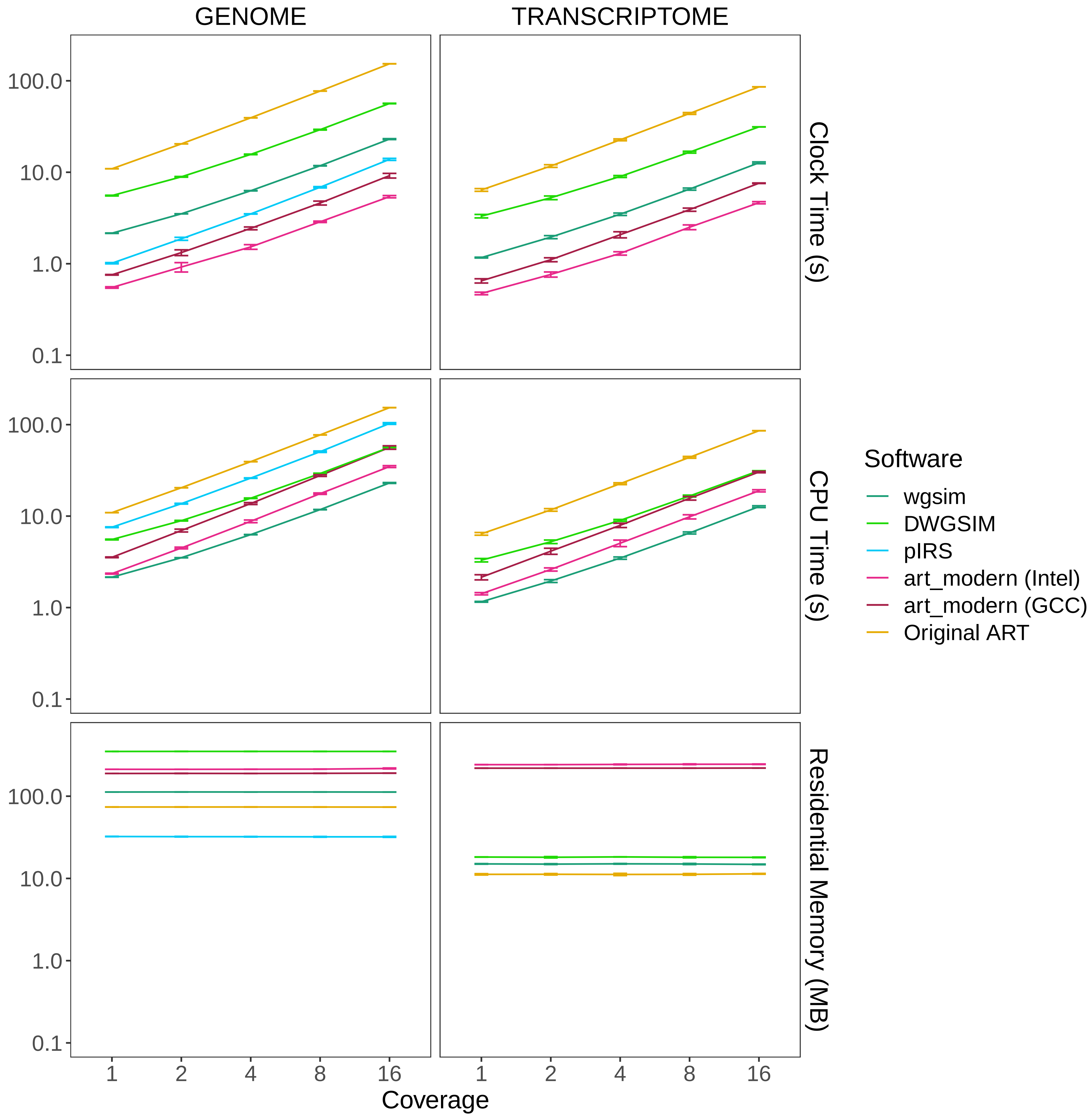

### time_memory_300.pdf

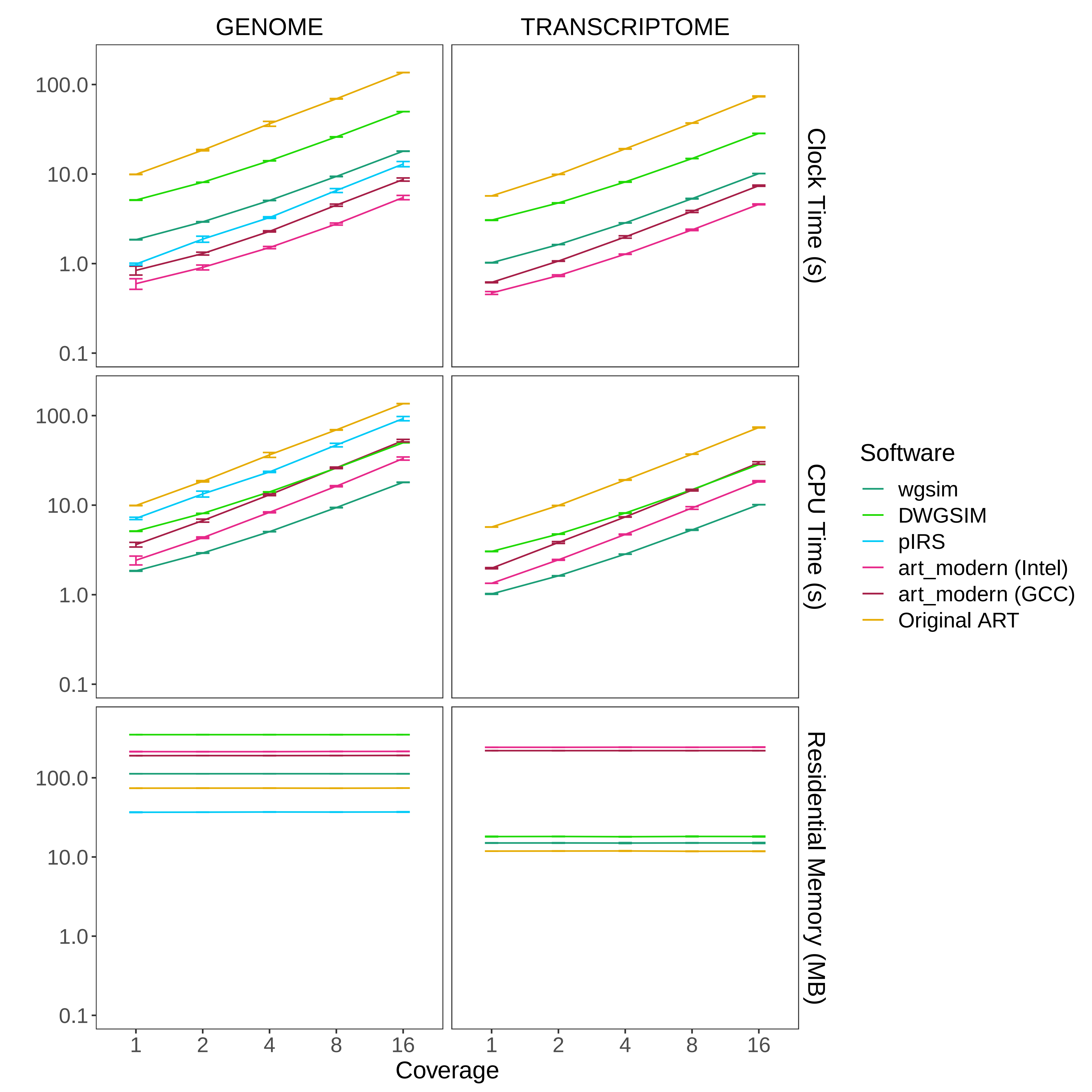
